# Altered dendritic spine function and integration in a mouse model of Fragile X Syndrome

**DOI:** 10.1101/396986

**Authors:** Sam A. Booker, Aleksander P.F. Domanski, Owen R. Dando, Adam D. Jackson, John T.R. Isaac, Giles E. Hardingham, David J.A. Wyllie, Peter C. Kind

## Abstract

Cellular and circuit hyperexcitability are core features of Fragile X Syndrome and related autism spectrum disorder models. However, a synaptic basis for this hyperexcitability has proved elusive. We show in a mouse model of Fragile X Syndrome, glutamate uncaging onto individual dendritic spines yields stronger single-spine excitation than wild-type, with more silent spines. Furthermore, near-simultaneous uncaging at multiple spines revealed fewer spines are required to trigger an action potential. This arose, in part, from increased dendritic gain due to increased intrinsic excitability, resulting from reduced hyperpolarization-activated currents. Super-resolution microscopy revealed no change in dendritic spine morphology, pointing to an absence of a structure-function relationship. However, ultrastructural analysis revealed a 3-fold increase in multiply-innervated spines, accounting for the increased single-spine excitatory currents following glutamate uncaging. Thus, loss of FMRP causes abnormal synaptogenesis, leading to large numbers of poly-synaptic spines despite normal spine morphology, thus explaining the synaptic perturbations underlying circuit hyperexcitability.

---

Cell and circuit hyperexcitability have long been hypothesized to underlie many core symptoms of Fragile X Syndrome (FXS) and autism spectrum disorders more generally, which includes sensory hypersensitivity, seizures and irritability (Contractor, Klyachko, and Portera-Cailliau 2015). The fundamental role of cellular excitability in circuit function raises the possibility that an alterations in neuronal intrinsic physiology may underlie a range of functional endophenotypes in FXS. Despite this potential link, few studies have examined the combined synaptic, dendritic, and cellular mechanisms that lead to generation of neuronal hyperexcitability during early postnatal development.

Many cellular properties are known to regulate neuronal excitability, ranging from neuronal morphology and intrinsic physiology, to synaptic transmission and plasticity. In FXS, a central hypothesis is that glutamatergic signalling at dendritic spines is impaired (Pfeiffer and Huber 2007, Irwin, Galvez, and Greenough 2000). The first major alteration described was a change in dendritic spine density and morphology (Irwin, Galvez, and Greenough 2000, Comery et al. 1997), however this observation was not apparent when examined at the nanoscale using super-resolution imaging methods (Wijetunge et al. 2014). Furthermore, despite these reported changes in spine density, no study has observed a change in synaptic event frequency that would be predicted by a change in spine density. This has important implications for our understanding of the synaptic aetiology of FXS, as many of the current theories are reliant on altered synaptic function (Bear, Huber, and Warren 2004, Pfeiffer and Huber 2009).

The rodent somatosensory cortex (S1) is well characterised in terms of its processing of tactile inputs, which, in the case of the barrel cortex arise from the whiskers on the facepads via relay synapses in the brainstem and ventrobasal thalamus (Schubert, Kötter, and Staiger 2007). The thalamic inputs arrive predominantly onto layer 4 stellate cells (L4 SCs) which integrate this information within L4, then project to L2/3 and L6. Furthermore, L4 SCs undergo a well described critical period for synaptic plasticity, which closes at postnatal day 7-8 (P7-8). For these reasons, L4 of S1 provides a well-described reductionist system to examine sensory processing (Fox 1992, Petersen 2007). Indeed, hyperexcitability has been observed within S1 of *Fmr1*^*-/y*^ mice, due in part to changes in intrinsic neuronal excitability, axonal morphology, and synaptic connectivity, which together result in increased network excitability (Bureau, Shepherd, and Svoboda 2008, Gibson et al. 2008, Zhang et al. 2014). The finding that the developmental critical period for thalamocortical synaptic plasticity is delayed in *Fmr1*^*-/y*^ mice relative to controls gave a suggestion as to how these cellular deficits may arise (Harlow et al. 2010). However, the effect of this delay in synaptic development affected dendritic spine function is not known. Furthermore, no study has directly examined how dendrites integrate synaptic inputs in the absence of FMRP, despite the fact that dendritic integration plays a key role in regulating cellular excitability (Magee 1999, Branco and Häusser 2011, Lavzin et al. 2012). In this context, the fact that intrinsic physiological deficits potentially associated with altered integration, particularly in HCN channel expression is of particular relevance (Gibson et al. 2008, Brager, Akhavan, and Johnston 2012, Zhang et al. 2014). Here, we directly test whether there is a functional relationship between dendritic spine function, intrinsic neuronal physiology and the role of HCN channels, dendritic integration, and ultimately neuronal output. To address this question, we use an integrative approach that combines whole-cell patch-clamp recording from neurons in S1 at P10-14 with 2-photon glutamate uncaging, *post-hoc* stimulated emission-depletion (STED) microscopy, and serial block-face scanning electron microscopy.

## Results

To assess the function of identified dendritic spines in *Fmr1*^-/y^ mice, we first performed single spine 2-photon glutamate uncaging. Whole-cell patch-clamp recordings were performed from layer 4 spiny-stellate cells (L4 SCs) in voltage-clamp, with a Cs-gluconate based intracellular solution containing a fluorescent dye (Alexafluor 488, 100 µM) and biocytin to allow on-line and *post-hoc* visualization of dendritic spines. Following filling we performed 2-photon photolysis of caged-glutamate (Rubi-glutamate) to elicit uncaged excitatory post-synaptic currents (uEPSCs; **Figure 1A**). From both the concentration- and power-response relationships, we determined that 300 µM [Rubi-Glutamate] and 80-100 mW laser power (λ780 nm) were optimal to produce saturating uEPSCs at -70 mV, which were fully abolished by CNQX, confirming that they were mediated by AMPA receptors (AMPARs). Analysis of the spatial properties of Rubi-Glutamate uncaging confirmed that the optimal position for photolysis was 0-1 µm from the edge of the spine head (**Figure S1**).

**Figure 1:**
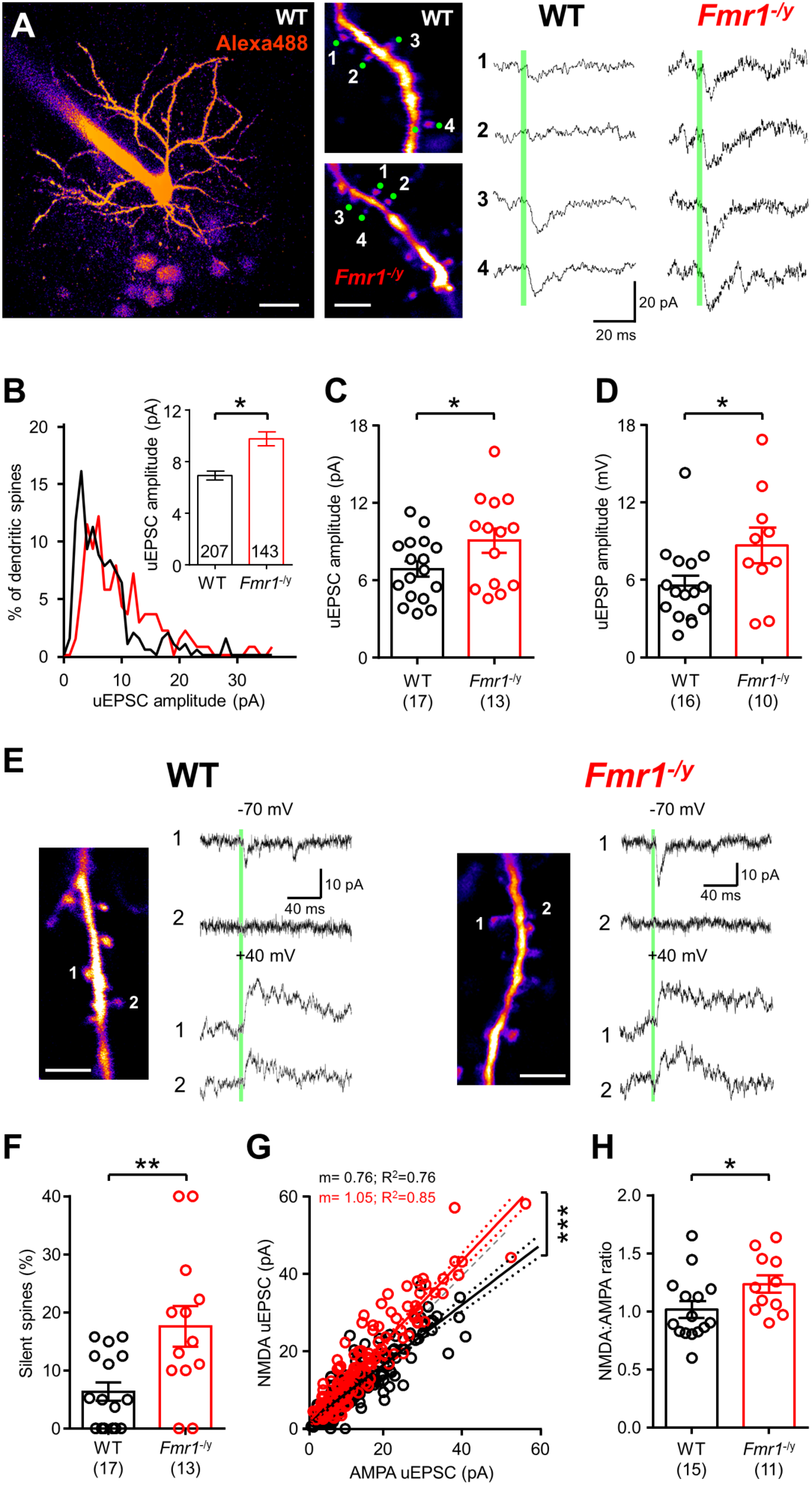
Dendritic spines on L4 SCs have larger uEPSCs with more silent synapses in *Fmr1*-/y mice. **A** 2-photon image of a L4 SC (left) with selected spines and AMPAR uEPSCs from WT and *Fmr1*-/y mice. Scale bars: 20 µm (left), 5 µm (right). **B** Single spine uEPSCs from WT (black) and *Fmr1*-/y (red) mice shown as a histogram, with spine average shown (inset). **C** Animal average uEPSC amplitudes, number of animals tested shown in parenthesis. **D** Animal average of uEPSP amplitudes. **E** AMPAR (upper) and NMDAR (lower) uEPSCs, illustrating “silent spines”. Scale: 5 µm. **F** Incidence of “silent spines” in WT and *Fmr1*-/y mice. **G** AMPAR and NMDAR uEPSCs for all spines, with NMDA/AMPA ratio (WT: 0.76 ± 0.03; *Fmr1*-/y; 1.05 ± 0.04; d.f.: 1, 331; F = 37.4; *p* <0.0001; F-test). **H** Animal averaged NMDA/AMPA ratio. Statistics shown: * - *p*<0.05, ** - *p*<0.01.

Comparison between genotypes revealed that the single spine uEPSCs in WT mice had an amplitude of 6.9 ± 0.4 pA (n=17 mice), while *Fmr1*^-/y^ mice (n=13 mice) showed a larger uEPSC amplitude of 9.8 ± 0.5 pA (d.f: 4, 5 χ^2^ = 8.26 *p* = 0.004; GLMM**, see Figure S2**), indicating that spines in *Fmr1*^-/y^ mice are enriched for AMPAR-mediated currents (**Figure 1B, C**). This difference appeared to be due to a greater population of uEPSCs at *Fmr1*^-/y^ spines with amplitudes over 10 pA (**Figure 1B**). As expected from a larger underlying current, the single spine uncaging excitatory post-synaptic potential (uEPSP) was also larger in *Fmr1*^-/y^ mice (0.73 ± 0.12 mV, n = 10 mice), when compared to WT littermates (0.47 ± 0.06 mV, n = 16 mice; d.f.: 24; t = 2.09; *p* = 0.046; T-test; **Figure 1D**). In a subset of dendritic spines we observed no AMPAR current when recorded at -70 mV, however a large NMDA receptor (NMDAR) current was present at +40 mV, indicating the presence of “silent” dendritic spines (**Figure E**). Quantification of the incidence of “silent” spines revealed an incidence of 17.6 ± 3.5% in *Fmr1*^-/y^ mice (n=13 mice), almost 3-fold higher than in WT mice (6.4 ± 1.6%, n=17 mice; d.f.: 27; t = 3.1; *p* = 0.005; T-test). When measured across all spines, the NMDA/AMPA ratio was significantly elevated as both a spine average (**Figure 1G**) and also as an animal average (Figure 1H), with *Fmr1*^-/y^ mice having a ratio of 1.24 ± 0.08 (n=11 mice) and WT of 1.02 ± 0.07 (n=15 mice; d.f: 4, 5 χ^2^ = 6.27 *p* = 0.012, GLMM, see also Figure S3).

Given that majority dendritic spines on L4 SC dendrites are formed by cortico-cortical synapses in WT mice (White and Rock 1980) and therefore likely constitute the majority of uncaged spines, we next asked whether synapses formed between L4 SCs had larger EPSC amplitudes by performing paired recordings between synaptically coupled neurons (**Figure 2**). As previously described in 2-week old mice (Gibson et al. 2008), we observed a reduced connectivity between L4 SCs in *Fmr1*^-/y^ mice, with a probability of only 14.8%, 50% lower than that of WT mice which had a connectivity of 33.6% (*p* = 0.015, Fisher’s exact test**, Figure 2C**). Despite this reduced connectivity, there was no difference in either failure rate (d.f.: 41; U = 129; *p* = 0.74; Mann-Whitney U test; **Figure 2D**) or unitary EPSC amplitude (d.f.: 51; U = 228; *p* = 0.14; Mann-Whitney U test, **Figure 2E**), suggesting that synaptic strength is unchanged at the majority of unitary synaptic contacts in *Fmr1*^-/y^ mice.

**Figure 2:**
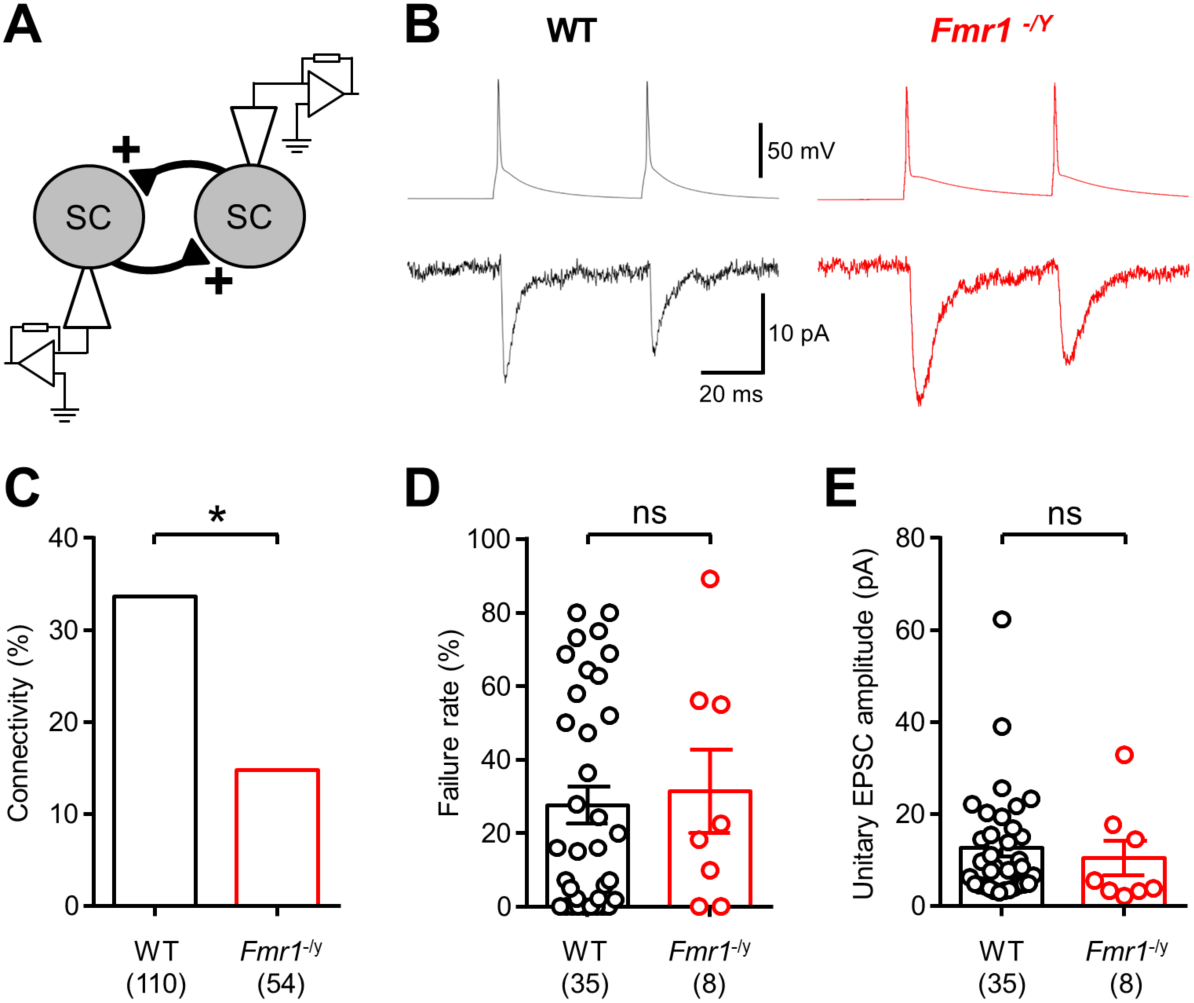
Typical EPSC amplitude at unitary connections between L4 SCs. **A** Schematic paired recordings between synaptically coupled L4 SCs. **B** representative presynaptic action potentials (top) produced unitary EPSCs in the second L4 SC (lower), from WT (black) and *Fmr1*-/y (red) mice. **C** Synaptic connectivity is reduced between L4 SCs in the *Fmr1*-/y mouse (d.f.: 162; *p* = 0.015; Fisher’s exact test); number of tested pairs is indicated. **D** Failure rate was not different between genotypes when a connection was present (d.f.: 41; U = 129; *p* = 0.74; Mann-Whitney U test). **E** Unitary EPSC amplitudes from L4 SC synapses were not different (d.f.: 41; U = 104; *p* = 0.22; Mann-Whitney U test), nor were WT unitary EPSCs different to *Fmr1*^-/y^ unitary EPSCs (d.f.: 51; U = 228; *p* = 0.14; Mann-Whitney U test). Statistics shown: ns – *p*>0.05, * - *p*<0.05.

The inclusion of biocytin within the recording solution allowed *post-hoc* visualisation of the recorded neurons, following fixation and re-sectioning. We next performed correlated Stimulated Emission-Depletion (STED) imaging of the same dendritic spines we had uncaged upon (**Figure 3A-E**). Measurement of nanoscale spine morphology revealed that there was no difference in either spine head width (**Figure 3B),** nor neck-length (**Figure 3D**), between WT (n=6 mice) and *Fmr1*^-/y^ (n=4 mice) mice.

**Figure 3:**
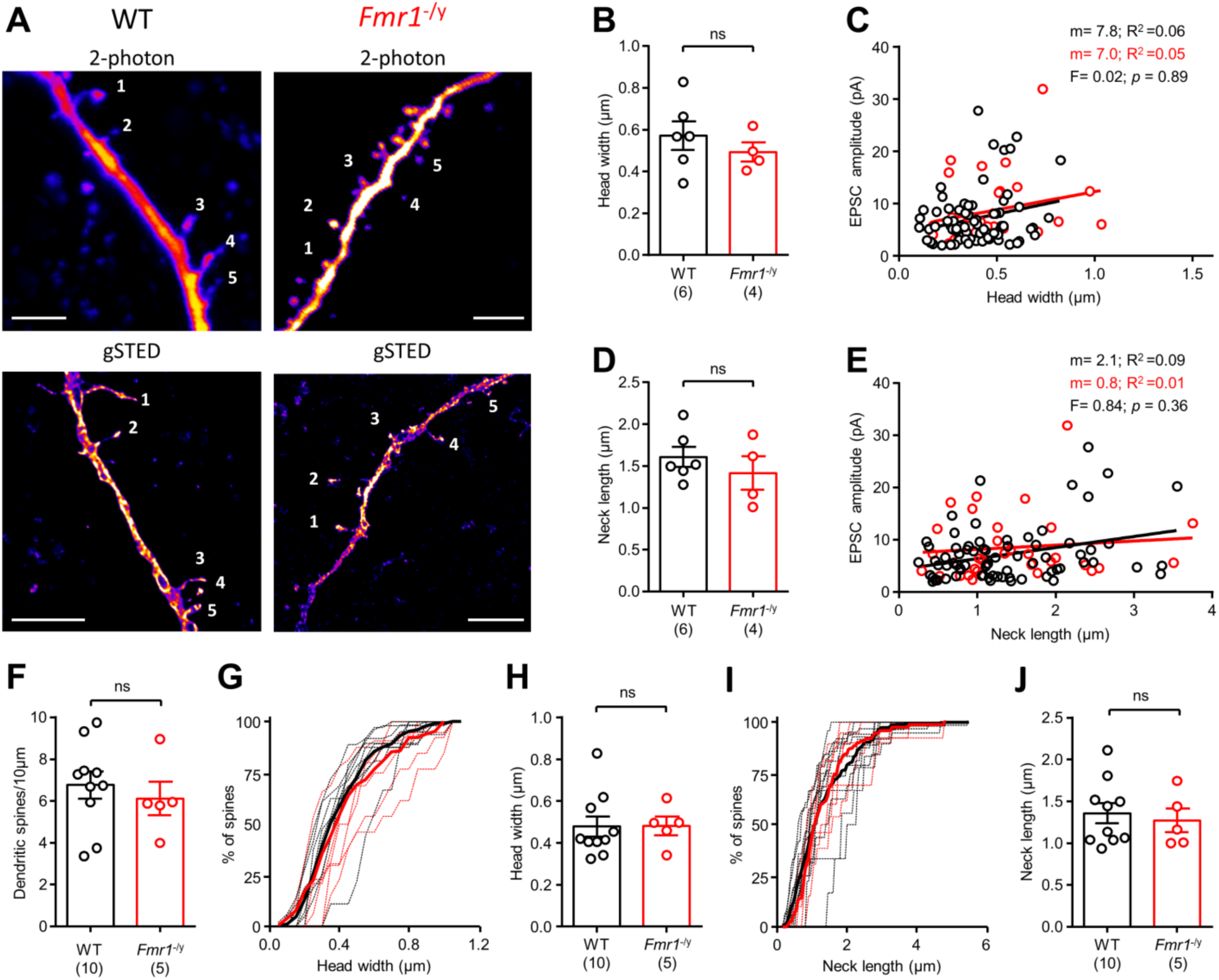
Dendritic spines show no difference in nanoscale morphology, or structure-function relationship. **A** Dendrites from WT (left) and *Fmr1*-/y (right) mice under 2-photon microscopy (top), then *post-hoc* STED imaging (bottom). Scale bar: 5 µm. **B** Average spine head width in WT (black) and *Fmr1*-/y (red) mice (WT: 0.43 ± 0.05; *Fmr1*-/y; 0.45 ± 0.04; d.f.: 8; t = 0.29; p = 0.78, T-test). Number of mice is indicated. **C** Comparison of spine head-width and uEPSC amplitude (comparing slope: d.f.: 1, 100; F = 0.02; p = 0.89). WT spines showed a positive correlation (d.f. 70, F=4.27, *p* = 0.042, F-test). **D** Average spine neck length (WT: 1.52 ± 0.22; *Fmr1*-/y; 1.31 ± 0.20; d.f.: 8; t = 0.66; p = 0.53, T-test; F-test). **E** Comparison of spine neck-width and uEPSC amplitude (Slope: WT: 2.1 ± 0.8; Fmr1-/y; 0.8 ± 1.4; d.f.: 1, 101; F = 0.84; p = 0.36; F-test). **F** Spine density on L4 SCs (WT: 6.8 ± 0.7 spines/10 µm; *Fmr1*-/y: 6.1 ± 0.80 spines /10 µm; d.f.: 13; t = 0.60; *p* = 0.56; T-test). **G** Distribution of non-uncaged spine head-widths, as an average of all mice (bold) and individual mice (dashed). **H** Average head-width of non-uncaged spines (WT: 0.48 ± 0.05 µm; *Fmr1*-/y: 0.48 ± 0.04 µm; d.f.: 13; U = 20.0; *p* = 0.59; Mann-Whitney U-test). **I** Distribution of spine neck-length of non-uncaged spines. **J** Average of spine neck-length in non-uncaged spines (WT: 1.36 ± 0.12 µm; *Fmr1*-/y: 1.27 ± 0.14 µm; d.f.: 13; U = 20.0; *p* = 0.55; Mann-Whitney U-test). Statistics shown: ns – *p* > 0.05 T-test.

Consistent with earlier findings (Ashby and Isaac 2011), we observed a weak positive correlation with spine head width and EPSC amplitude in WT mice (7.8 ± 3.8 pA/µm, R^2^=0.06, F=4.3, *p*=0.042, F-test), which was not different to that of *Fmr1*^*-/y*^ mice (F=0.02, *p*=0.89, Sum-of-Squares F-test; **Figure 3C**). We observed no correlation with spine neck length and EPSC amplitude (**Figure 3E**). To confirm that uncaging itself did not have an effect on any potential difference, we also measured spines from non-uncaged dendrites on filled neurons; spine density was not different between genotypes (**Figure 3F**). Likewise head width (**Figure 3G, H**) and neck length (**Figure 3I, J**) did not show any statistical difference, in agreement with previous findings in L5 of S1 and CA1 of the hippocampus (Wijetunge et al. 2014).

Given the synaptic strengthening of individual dendritic spines, but the lack of change in unitary EPSC amplitudes and spine morphology, we next asked whether the synaptic structure of dendritic spines was altered. To achieve this we used serial block-face scanning electron microscopy in L4 of S1 from mice perfusion fixed at P14. In serial stacks (50 nm sections; **Figure 4**) we identified Type-1 asymmetric synapses on dendritic spines, based on the presence an electron dense post-synaptic density (PSD) opposing an axon bouton containg round vesicles. Following 3-dimensional reconstruction, we identified a subset of dendritic spines that contained more than 1 PSD, which were each contacted by an independent presynaptic axon bouton (**Figure 4A, B**), and henceforth referred to as multi-innervated spines (MIS). These MIS were present in both genotypes, however the incidence in *Fmr1*^-/y^ mice was 20.5 ± 1.6% of all spines (n=7 mice), 3-fold higher than in WT littermates (7.2 ± 1.5% of spines, n=3 mice, d.f.: 8; t = 4.9; *p* = 0.001; T-test; **Figure 4C**), which was similar to previously observed levels in organotypic slice cultures from WT mouse hippocampus (Nikonenko, Jourdain, and Muller 2003).

**Figure 4:**
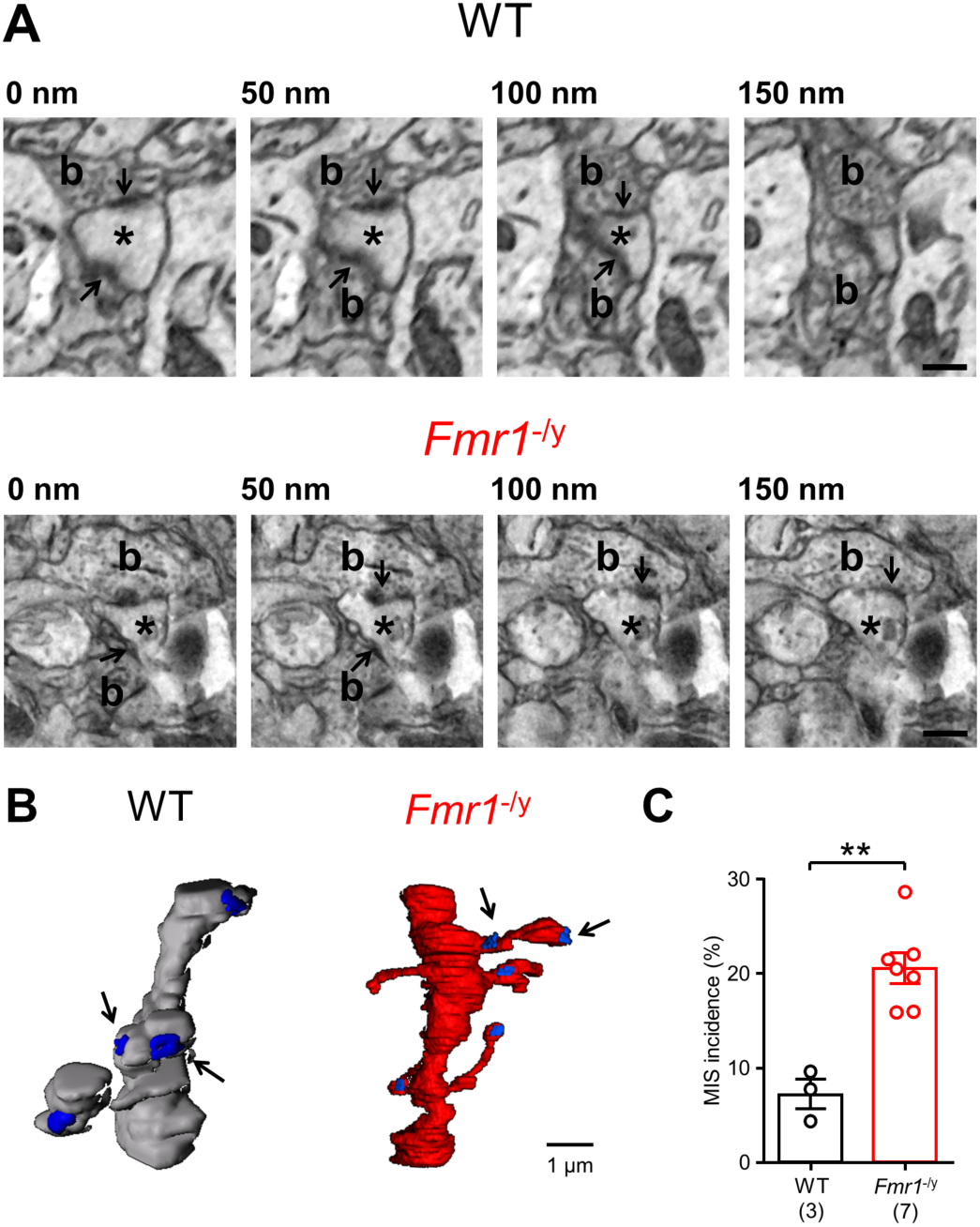
L4 spines in *Fmr1*^-/y^ mice form multiple synaptic contacts. **A** Serial electron micrographs in L4 from WT and *Fmr1*^-/y^ mice, indicating spines (asterisk) contacted by multiple presynaptic boutons (b) each with a PSD (arrows); scale bar: 500 nm. **B** Reconstructed dendrites from WT (grey) and *Fmr1*^-/y^ (red) mice, with PSDs (blue) and MIS indicated (arrows). **C** Incidence of MIS in WT and *Fmr1*^-/y^ mice. Statistics shown: ** - *p*<0.01.

The presence of higher numbers of MIS in *Fmr1*^-/y^ mice, and larger single spines uEPSCs, despite a similar density of spines and similar dendritic morphologies (Till et al. 2012), would suggest an increased number of synapses for each L4 SC. The conventional method to assess such a change in synapse number is to perform miniature EPSC (mEPSC) recordings (**Figure 5A**). AMPAR mEPSCs recorded at -70 mV in *Fmr1*^-/y^ mice were very similar to WT in both amplitude (d.f.: 47; t = 0.25; *p* = 0.81; T-test) and frequency (d.f.: 47; t = 1.3; *p* = 0.19; T-test; **Figure 5B**). NMDAR mEPSCs, recorded at +40 mV in the presence of CNQX, also had very similar amplitudes (d.f.: 17; t = 0.85; *p* = 0.41; T-test). However, *Fmr1*^-/y^ mice showed a 54% increase in NMDAR mEPSC frequency compared to WT mice (d.f.: 17; t = 2.4; *p* = 0.03; T-test; **Figure 5C**). These data indicate that while AMPAR-containing synapses number and strength are unaltered in *Fmr1*^-/y^ mice, they possess ∼50% more NMDAR containing synapses.

**Figure 5:**
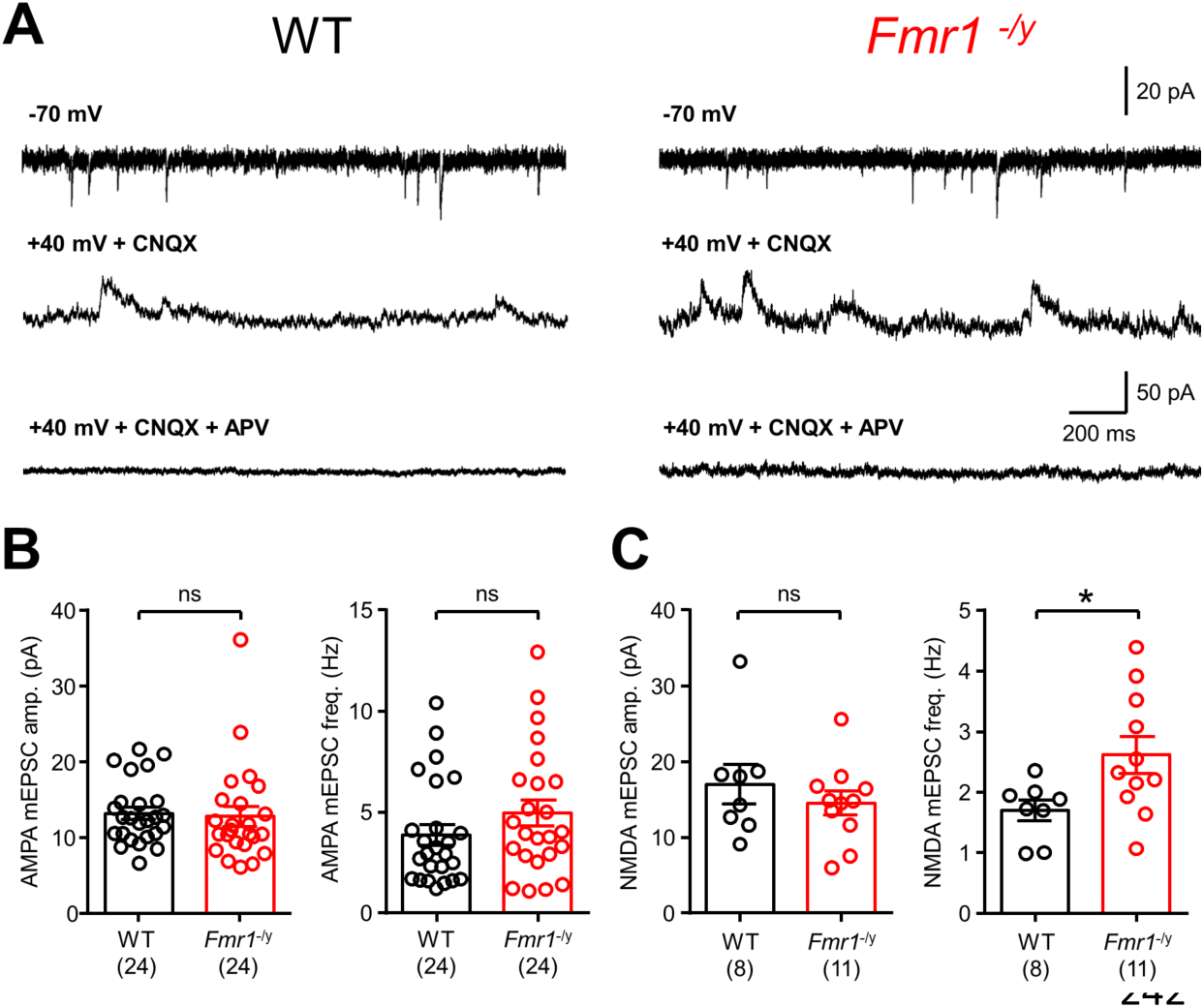
mEPSCs in *Fmr1*-/y L4 SCs show enrichment of NMDAR synapses. **A** mEPSCs recorded from L4 SCs for AMPAR at -70 mV (top), NMDAR at +40 mV with CNQX (10 µM, middle), and following application of the NMDAR antagonist D-AP5 (50 µM, bottom) in the same cell; from WT (left) and *Fmr1*-/y (right) mice. **B** Quantification of AMPAR mEPSC amplitude (WT: 13.1 ± 0.8; *Fmr1*-/y; 12.7 ± 1.3) and frequency (WT: 3.9 ± 0.5; *Fmr1*-/y; 4.9 ± 0.6;) in WT (black) and *Fmr1*-/y (red) mice. Number of mice indicated in parenthesis. **C** NMDAR mEPSC amplitude (WT: 16.9 ± 2.6; *Fmr1*-/y; 14.4 ± 1.6; and frequency (WT: 1.7 ± 0.17; *Fmr1*-/y; 2.6 ± 0.3;) measured in WT and *Fmr1*^-/y^ mice. Statistics shown: ns – *p*>0.05, * - *p*<0.05. All data shown as mean ± SEM.

While these observed changes in synaptic properties reveal differences in dendritic spine function, alone they do not reveal how neurons integrate excitatory inputs leading to hyperexcitability. Dendritic spines act as spatiotemporal filters whose summation is dependent upon the cable properties of dendrites (Rall 1959), synaptic receptor content (Lavzin et al. 2012) and the intrinsic membrane properties (Magee and Johnston 1995, Branco and Häusser 2011).

Therefore, we next measured intrinsic excitability of L4 SCs in response to hyper- to depolarising current injections (**Figure 6A, B**). Input resistance, as measured from the steady-state current-voltage relationship (**Figure 6C**) and smallest current step response (Figure 6C, inset) was increased in *Fmr1*^-/y^ mice, compared to WT (d.f.: 108; t = 2.24; *p* = 0.027; T-test). This led to an increase in action potential (AP) discharge in *Fmr1*^-/y^ mice (**Figure 6D**, d.f.: 5, 633; F = 4.89; *p* = 0.0002; 2-way ANOVA), resulting from a decreased rheobase currents in the recorded L4 SCs (Figure 6D, inset; d.f.: 108; t = 2.28; *p* = 0.025; T-test). This increase in input resistance was also matched by an increase in impedance, when measured with a sinusoidal wave (0.2 – 20 Hz, 50 pA, 20 s duration, **Figure 6E**), which was associated with a resonant frequency of 1.1 ± 0.1 Hz in L4 SCs from *Fmr1*^-/y^ mice, higher than that of 0.8 ± 0.1 Hz in WT littermates (d.f.: 25; t = 3.2; *p* = 0.004; T-test; **Figure 6F**); this was not matched by a change in resonant dampening (Q-factor: WT: 1.23 ± 0.07; *Fmr1*^-/y^; 1.13 ± 0.03; d.f.: 24; t = 0.7; *p* = 0.49; T-test) suggesting equally sustained activity at these resonance frequencies in both genotypes. This analysis demonstrates that L4 SCs from *Fmr1*^-/y^ mice are intrinsically more excitable and tune to higher frequencies than their WT counterparts.

**Figure 6:**
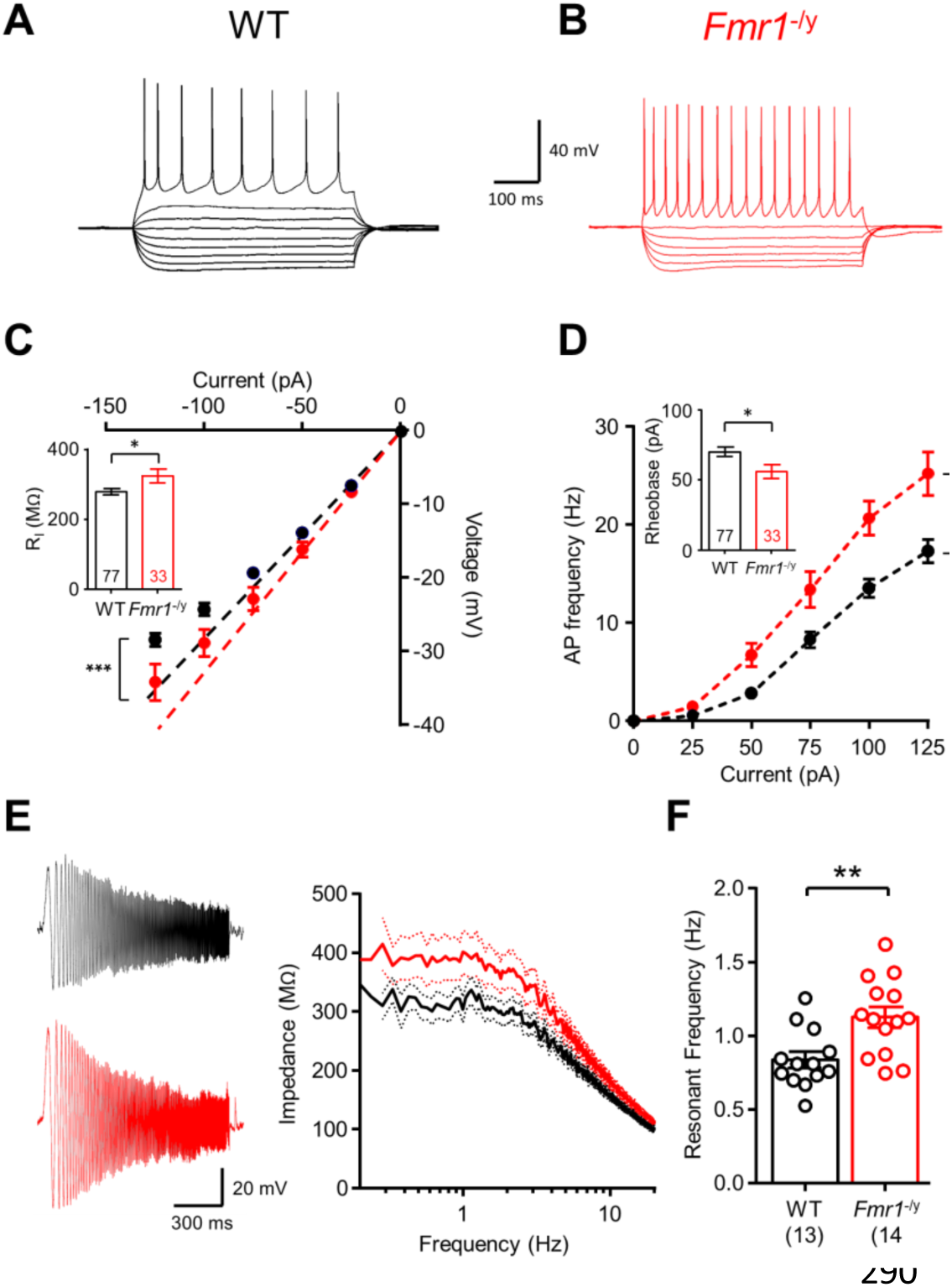
Altered intrinsic physiological properties of L4 SCs in *Fmr1*-/y mice resulting from decreased I_h_. Voltage responses to hyper- to depolarizing currents (−125 to +125 pA, 25 pA steps, 500 ms duration) results in action potentials (AP) in WT (**A**) and *Fmr1*-/y (**B**) mice. **C** The current-voltage response to hyperpolarizing currents with linear fit (dashed lines) in WT (black) and *Fmr1*-/y (red) mice (d.f.: 1, 326; F = 4.79; *p* = 0.030; F-test). **C, inset** Input resistance (R_I_) measured from all L4 SCs tested. **D** Current-frequency plot showing AP discharge. **D, inset** Average rheobase current measured in all cells. **E**, subthreshold membrane chirps (0.2 -20 Hz, 50 pA, 20 s duration) in L4 SCs from WT (black) and *Fmr1*-/y mice. Right, frequency-impedance plot for both genotypes ± SEM, shown on a logarithmic frequency scale. **F**, measured resonant frequency of L4 SCs from both genotypes. Statistics shown: * - *p* < 0.05, ** - p < 0.01.

It has been shown in S1 L5 pyramidal cells that HCN channels producing the hyperpolarisation-activated current (I_h_) are reduced when measured indirectly as a voltage “sag” in current-clamp (Zhang et al. 2014). Therefore, we next asked to what level I_h_ is present in L4 SCs and what its effect is on their excitability. To address this we directly measured I_h_ in voltage clamp (−60 mV holding potential) in response to hyperpolarising voltage steps (0 to -50 mV, 5 mV steps, 500 ms duration).

WT L4 SCs possessed large I_h_ at hyperpolarised potentials with a maximum amplitude of 25.0 ± 2.9 mV, whereas *Fmr1*^-/y^ L4 SCs showed reduced I_h_ activation overall (d.f.: 10, 310; F = 3.94; *p* < 0.0001 for interaction; 2-way ANOVA; **Figure 7A**), which had a peak amplitude of 15.7 ± 1.7 mV, 30% lower than WT cells (*p* = 0.003, Bonferroni post-test). We next applied the I_h_ blocker ZD-7,288 (ZD; 20 µM) to a subset of cells to assess the effect of I_h_ on cellular excitability. In these cells, we confirmed increased baseline input resistances in *Fmr1*^-/y^ mice compared to WT (d.f.: 44; t = 2.18; *p* = 0.034; T-test; **Figure 7B**), which in WT cells was increased by 61% following ZD application (d.f.: 21; t = 8.30; *p* < 0.0001; Paired T-test), while *Fmr1*^-/y^ L4 SCs showed a 30% increase (d.f.: 20; t = 3.57; *p* = 0.002; Paired T-test), 2-fold lower than WT cells (d.f.: 58; t = 2.75; *p* = 0.008; T-test; **Figure 7C**). Given the observed differences in AP discharge (**Figure 6D**), we next tested whether ZD normalised firing properties. In WT L4 SCs, ZD application significantly increased AP firing (d.f.: 5, 155; F = 7.5; *p* < 0.0001 for interaction; 2-way ANOVA; **Figure 7D**), but did not alter the peak firing (p = 0.31, Bonferroni post-test), meanwhile ZD had no effect on the AP discharge of *Fmr1*^-/y^ L4 SCs (d.f.: 5, 174; F = 0.23; *p* = 0.95 for interaction; 2-way ANOVA; **Figure 7E**). Finally, we asked what effect ZD had on the resonant properties of L4 SCs. In WT L4 SCs, ZD application strongly increased the impedance of WT neurons at low frequencies (**Figure 7F, H**), which lead to a 33% increase in impedance at the resonant frequency (d.f.: 6; t = 3.81; *p* = 0.009; Paired T-test), whereas ZD had no effect on impedance in *Fmr1*^-/y^ neurons (d.f.: 7; t = 1.61; *p* = 0.15; Paired T-test; **Figure 7G, H**). Together these data show that the intrinsic excitability of L4 SCs is increased in *Fmr1*^-/y^ mice, due to reduced I_h_ in them, which itself can fully explain genotype specific differences in cellular intrinsic excitability.

**Figure 7:**
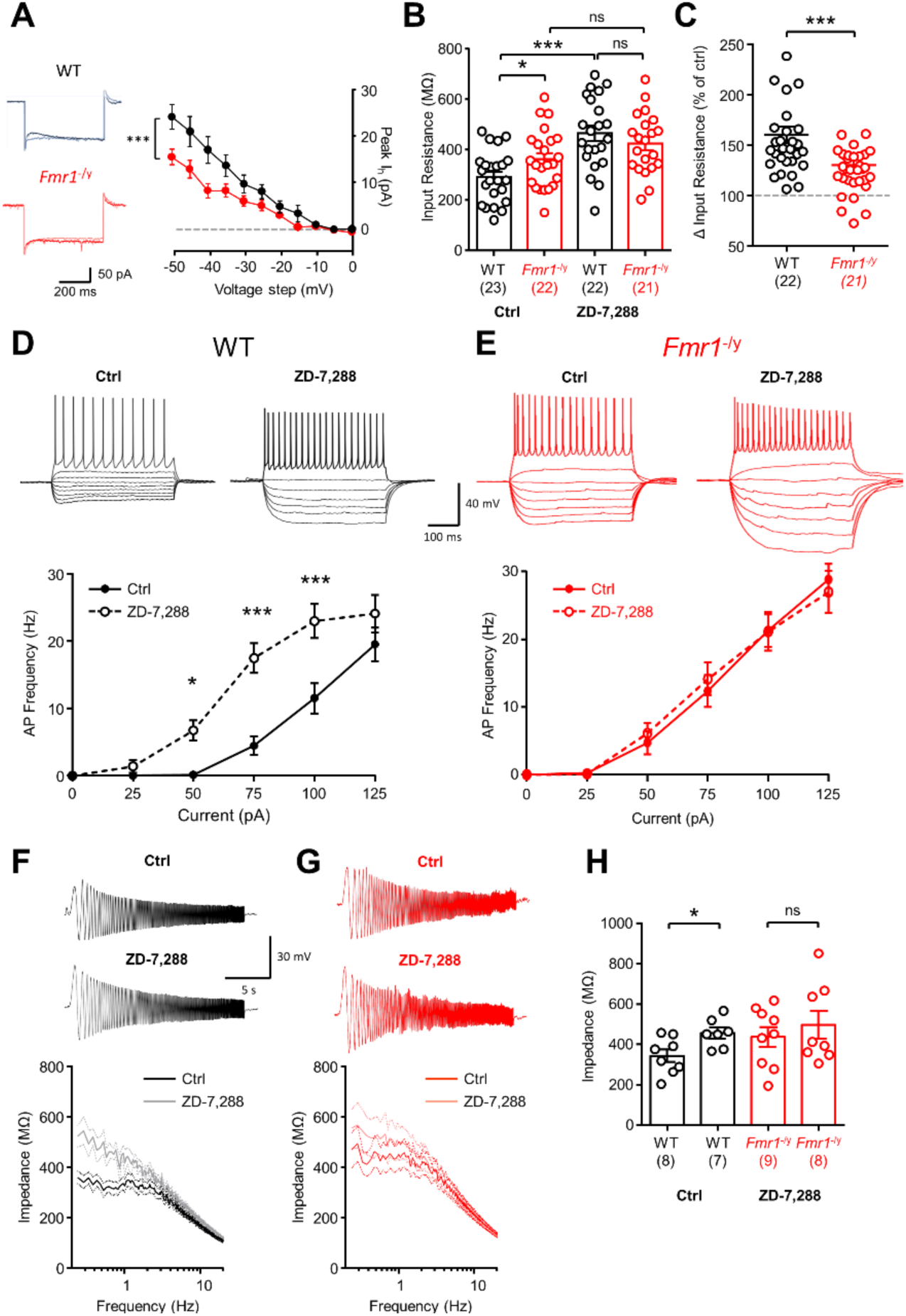
HCN channel mediated I_h_ is reduced in L4 SCs from *Fmr1*^-*/y*^ mice leading to enhanced excitability. **A** left, Hyperpolarizing steps in L4 SCs at -60 mV resulting in I_h_. Right, measured I_h_ amplitude at all tested hyperpolarising steps. **B** R_I_ measured under control conditions and following bath application of the I_h_ blocker ZD-7,288 (ZD; 20 µM) (WT: d.f.: 5; t = 7.19; *p* = 0.0008; paired T-test; *Fmr1*^-/y^: d.f.: 5; t = 4.52; *p* = 0.0063; paired T-test). **C**, measured change in input resistance change following ZD application (as 100% of control levels). **D,** hyper-to depolarising current steps (− 125 to +125 pA, 25 pA steps, 500 ms duration) in WT L4 SCs under control conditions and following ZD-7,288 application. Lower, Current-frequency plot of AP discharge frequency before (solid lines) and after (dashed lines) ZD application. **E**, current-frequency responses according to the same scheme as C, but in *Fmr1*^-/y^ L4 SCs. **F**, subthreshold membrane chirps (0.2-20 Hz, 50 pA, 20 s duration) and current-impedance plot for L4 SCs from WT mice before (black) and after (grey) ZD application. **G**, The same data as in F, but in *Fmr1*^-/y^ mice. **H**, Impedance measured at peak resonant frequency in WT and *Fmr1*^-/y^ L4 SCs before (Ctrl) and after ZD application. Statistics shown: ns – *p* >0.05 * - *p* < 0.05, *** - p < 0.001.

Given that I_h_ is a key determinant of dendritic integration (Magee 1999), we next wanted to assess both spatial and temporal dendritic summation in the *Fmr1*^-/y^ L4 SCs. To address spatial summation in L4 SC dendrites we performed near-simultaneous glutamate uncaging at multiple spines (**Fig. 8A**), by focal puff application of rubi-glutamate (10 mM) and rapidly uncaging on dendritic spines (0.5 ms/spine). We first performed a sequential uncaging (i.e. each spine individually), then near simultaneous uncaging of spine ensembles (i.e. groups of spines; **Figure 8B**). Summating EPSCs ultimately resulted in a AP discharge from L4 SCs. *Fmr1*^-/y^ L4 SCs required fewer spines on average to produce an AP (d.f.: 23; t = 2.3; *p* = 0.03, T-test; **Figure 8C**), which was more pronounced when silent spines were excluded from analysis (d.f.: 18; t = 3.2; *p* = 0.005). Measurement of the summated EPSP, with respect to number of spines near-simultaneously uncaged showed that both WT and *Fmr1*^-/y^ L4 SC dendrites had a linear increase in EPSP amplitude with increasing number of spines (**Figure 8D**), which was significantly greater in the *Fmr1*^-/y^ L4 SCs (d.f.: 1, 170; F = 8.98; *p* = 0.003; F-test). This measure, however, includes effects due to increased spine synaptic strength and input resistance, as well as dendritic integrative properties. Therefore, we next compared the expected linear sum of single spine EPSPs to that of the observed summated EPSP (**Figure 8E**), thereby excluding individual spine strength and input resistance effects on EPSP amplitude. We observed linear integration in WT and *Fmr1*^-/y^ L4 SCs, however WT neurons showed low levels of integration (Slope: 0.50 ± 0.09), while *Fmr1*^-/y^ neurons presented over 50% higher integration (Slope: 0.79 ± 0.08; d.f.: 1, 195; F = 3.18; *p* = 0.044; F-test). These data clearly show that the dendrites of *Fmr1*^-/y^ L4 SCs undergo excessive dendritic summation of synaptic inputs. To confirm that dendritic summation is altered in response to endogenous synaptic transmission, we next provided extracellular stimulation to thalamocortical afferents (TCA) from the ventrobasal thalamus, whilst recording from L4 SCs (**Figure 8F**). Stimulus intensity was titrated so that an EPSC of ∼150 pA was produced, then trains of EPSPs were elicited in current-clamp at either 5 or 10 Hz. At these stimulation intensities summating EPSPs in L4 SCs in WT mice never produced a somatic AP, however in *Fmr1*^-/y^ mice 5 Hz stimulation resulted in an AP in 19 ± 7% of recordings (d.f.: 16; t = 2.57 & 3.81; *p* = 0.02 & 0.002, T-test) and 10 Hz stimulation 55 ± 13% of the time (d.f.: 16; t = 3.81; *p* = 0.002, T-test), confirming that dendritic integration properties alter the output of L4 SCs, to promote hyperexcitability (**Figure 8G**).

**Figure 8:**
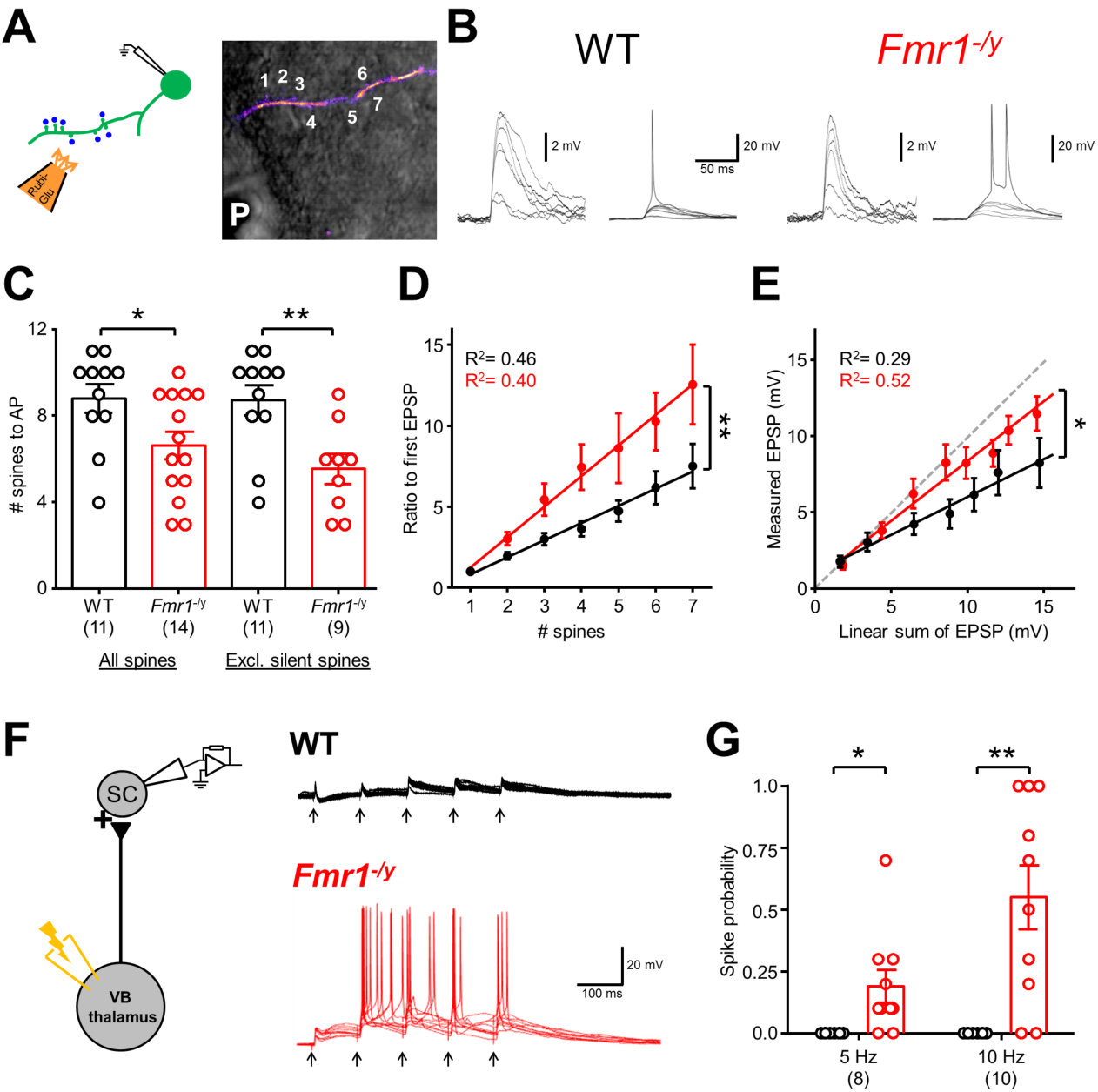
Enhanced dendritic integration of L4 SCs in *Fmr1*^-/y^ mice. **A** Schematic of near-simultaneous glutamate uncaging (Rubi-Glu) at multiple spines (blue dots/numbers). **B** Near-simultaneous glutamate uncaging produced subthreshold (inset, right) and suprathreshold uEPSPs (inset, left) along dendrites. **C** The number of spines required to evoke an AP, from all spines (left; WT: 8.8 ± 0.7; *Fmr1*^-/y^; 6.6 ± 0.6) and excluding “silent spines” (right; WT: 8.7 ± 0.7; *Fmr1*^-/y^; 5.6 ± 0.7). **D** Summation of near-simultaneous subthreshold uEPSPs normalized to the first EPSP in WT (black) and *Fmr1*^-/y^ (red) L4 SCs (Slope: WT: 1.1 ± 0.13; *Fmr1*^-/y^; 1.9 ± 0.2; d.f.: 1, 170; F = 8.98; *p* = 0.003; F-test). **E** Summating uEPSPs plotted against the expected linear-sum. Unity is indicated (grey). **F** Electrical stimulation of TCA at low frequency 10 Hz is shown. **G** Average spike probability in response to 5 Hz and 10 Hz stimulation. Statistics shown: * - *p*<0.05, ** - p<0.01.

I_h_ is a major factor contributing to the spatial and temporal summation of synaptic events, serving to shorten decay times of summating EPSPs (Magee 1999). Given the reduced I_h_ we observe in *Fmr1*^-/y^ L4 SCs, we next asked whether inhibition of the HCN channel resulted in altered decay time of integrated synaptic events. Summating uEPSPs from WT mice (normalised to the initial uEPSP) displayed long decay times at low summation, which were more rapid at higher summation levels (**Figure S4A, S4B**). By comparison, in *Fmr1*^-/y^ mice we did not observe these relationships and the genotype-specific slopes of the log(EPSP summation) were highly divergent (d.f.: 1, 109; *F* = 32.1, *p* <0.0001; F-test). The summation-dependent temporal sharpening of EPSPs in WT neurons was abolished following application of the I_h_ blocker ZD-7,288 (Comparing slope: d.f.: 1, 85; *F* = 6.4, *p*= 0.01; F-test; **Figure S4D**), and also prolonged decay times of the first EPSP (Figure 9F, d.f.: 15; t = 2.34; *p* = 0.034; T-test; **Figure S4C**). ZD-7,288 had no observable effect on summating EPSPs in *Fmr1*^-/y^ L4 SCs (**Figure S4E**). Taken together these data clearly show that dendrites of L4 SCs in the *Fmr1*^-/y^ undergo excessive summation, largely due to reduced I_h_ in these neurons.

## Discussion

L4 SCs in the primary somatosensory cortex are the first cortical cells to receive and integrate incoming sensory information, which is integrated and relayed within the cortex. As such, L4 SCs play a crucial role in sensory perception (Petersen 2007). Individuals with FXS show altered sensory processing (Lachiewicz et al. 1994, Miller et al. 1999) and mouse models show altered circuit processing in primary sensory areas (Bureau, Shepherd, and Svoboda 2008, Harlow et al. 2010, Deng, Sojka, and Klyachko 2011, Gonçalves et al. 2013, Zhang et al. 2014, Contractor, Klyachko, and Portera-Cailliau 2015). Furthermore, while FMRP has been shown repeatedly to regulate synapse function and plasticity, little is known about how these alterations affect dendritic spine function and dendritic integration to sensory input. To address these questions, we used glutamate uncaging at L4 SC dendritic spines to examine how they integrate and generate action potentials following synaptic stimulation. We show that L4 SCs in S1 have dendritic and synaptic properties that result in increased action potential generation in *Fmr1*^-/y^ mice relative to WT controls. Specifically, we show increased excitatory synaptic currents at individual spines resulting from increased AMPAR and NMDAR content. Despite this, we observed no change in spine morphology using STED microscopy and there was little correlation between spine structure and function, indicating that spine morphology is not an effective proxy for spine function, at least at the age used in this study. However, electron microscopic analysis revealed an increase in multiply-innervated spines which likely accounts for the increase in single-spine synaptic currents. Interestingly there was also an increase in silent spines which agrees with the increase in NMDAR mEPSC frequency, but not AMPAR mEPSC frequency. The overall increase in dendritic spine currents was accompanied by enhanced dendritic integration likely resulting, at least in part, from a ∼50% reduction in I_h_. This reduced I_h_ was causal to the altered intrinsic physiology of L4 SCs at P12-14. Finally, TCA stimulation at frequencies that fail to elicit AP discharge from L4 SCs in WT mice, in the presence of intact synaptic inhibition, reliably elicits APs in *Fmr1*^-/y^ neurons, indicating that the local inhibitory circuit cannot compensate for the increase in synaptic and dendritic excitability. Together these findings demonstrate that aberrant dendritic spine function and dendritic integration combine to result in cellular hyperexcitability in L4 SCs. As the first cortical cells to receive input from the sensory periphery, the resultant hyperexcitability likely contributes previously reported circuit excitability in *Fmr1*^*-/y*^ mice and the sensory hypersensitivities in individuals with FXS.

To our knowledge, this is the first study to quantify the incidence of MIS in intact tissue, and implicate their presence in pathological states associated with a disease model. Indeed, the mean increase in spine uEPSC amplitude in *Fmr1*^-/y^ mice is likely caused by the increase in the number of MIS. Indeed, the presence of MIS in both WT and *Fmr1*^-/y^ mice disagrees with the 1 spine/1 synapse hypothesis (Harris, Jensen, and Tsao 1992). A potential mechanistic link between loss of FMRP and the increase in MIS may come from its ability to regulate PSD-95. *Psd-95* mRNA is a known FMRP target (Darnell et al. 2011) and an increase in PSD-95 puncta in L4 of S1 has been observed (Wang, Smith, and Mourrain 2014), with no change in cell number, dendritic morphology, or spine density in *Fmr1*^-/y^ mice (Till et al. 2012). Furthermore, transient overexpression of PSD-95 results in increased MIS incidence through a nitric oxide synthase dependent mechanism (Nikonenko, Jourdain, and Muller 2003), which itself is another FMRP target (Darnell et al. 2011). Future experiments exploring the effect of NOS blockade or PSD-95 reduction in *Fmr1*^-/y^ mice would test this mechanistic relationship. Furthermore, how the presence of MIS influences dendritic protein synthesis and whether it presents a therapeutic target requires further study.

Interestingly the increase in spines with increased uEPSC amplitudes and MIS was mirrored by an increase in silent spines, though their number was insufficient to compensate for the overall increase in dendritic currents in other spines. An increase in silent TCA synapses at P7 (Harlow et al. 2010) was previously reported in *Fmr1*^-/y^ mice. However, this study also reported a delay in the critical period for inducing LTP at these synapses which terminated at P10. Therefore, the period of synaptic potentiation at TCA synapses is complete by the age we tested in this study. Hence the percentage of silent spines receiving TCA input would be expected to be low (Crocker-Buque et al. 2014). Furthermore, the reduced connectivity between L4 SCs at P12-14, despite no change in spine density (Till et al., 2012), strongly indicates that SC to SC synapses are preferentially silent at this developmental stage in the *Fmr1*^-/y^ mouse. Together, these findings suggest that silent spines measured in our study reflect cortico-cortical, rather than TCA, synapses. Given the hierarchical nature of sensory system development, it would not be surprising if a delay in intra-cortical synapse development in *Fmr1*^-/y^ mice follows the aforementioned delay in TCA synapse development, but this remains to be directly tested.

While dendritic spines are functionally disrupted in the *Fmr1*^-/y^ mouse, using super resolution microscopy we found no evidence of a genotypic difference in spine morphology of L4 SC neurons. This is in good agreement with our previous findings that spine morphology is unaffected in hippocampal CA1 and layer 5 S1 neurons (Wijetunge et al. 2014). Furthermore, we find only a weak correlation between dendritic spine structure and function, demonstrating the pitfalls of using spine structure as a proxy for synaptic function, especially in young animals and genetic models of disease. These findings are in stark contrast to those observed from post-mortem human tissue (Irwin, Galvez, and Greenough 2000) or from other mouse studies (Comery et al. 1997); however these studies were only performed with diffraction-limited microscopy, suggesting that super-resolution imaging techniques should be the gold-standard for dendritic spine morphological studies in future. Single dendritic spines do not typically produce AP discharge from neurons, rather they require co-activation and summation of multiple synaptic inputs arriving with high temporal precision (Losonczy and Magee 2006). L4 SCs have been previously been shown to possess linear integration of Ca^2+^ influx in their dendrites (Jia et al. 2014). We show that synaptic potentials also linearly integrate in L4 SCs of WT mice, and that this integration is strongly enhanced in *Fmr1*^-/y^, leading to more efficient discharge of APs, due in large part to a reduced I_h_. I_h_ has been implicated in the altered neuronal excitability of FXS (Brager, Akhavan, and Johnston 2012, Zhang et al. 2014), with the HCN1 channel expression dictating whether the current is increased or decreased. However, no study has until now shown a direct effect of I_h_ on whole-cell excitability. We see a strong reduction I_h_ in L4 SCs from *Fmr1*^-/y^ mice; thus inhibition of HCN channels alters intrinsic physiology and dendritic integration to a lesser degree than in WT mice. Given that I_h_ amplitude differs between regions and cell type, alteration in the density of HCN channels may reflect homeostatic regulation of neuronal excitability.

In summary, we provide the first direct evidence in *Fmr1*^-/y^ neurons for a functional deficit at excitatory synapses onto dendritic spines and that these alterations contribute to an increase in dendritic integration. The summation of synaptic responses contributes to hyperexcitability of sensory neurons in the *Fmr1*^-/y^ mouse, which along with changes in intrinsic excitability, may underlie pathophysiology associated with altered sensory function.

## Methods

### Animals and ethics

All procedures were performed in line with Home Office (ASPA, 2013; HO license: P1351480E) and institutional guidelines. All experiments were performed on C57/Bl6J mice, bred from *Fmr1*^+/-^ mothers, cross-bred with *Fmr1*^+/y^ male mice, giving a Mendelian 1:1 ratio of *Fmr1*^+/y^ and *Fmr1*^-/y^ amongst male offspring. Only male mice were used for the present study and all mice were killed at P10-15, before separation from the mother. Mothers were given *ad libitum* access to food and water and housed on a 12 hr light/dark cycle. All experiments and analysis were performed blind to genotype.

### Acute slice preparation

Acute brain slices were prepared similar to previously described (Agmon and Connors 1991, Booker, Song, and Vida 2014). Briefly, mice were decapitated without anaesthesia and the brain rapidly removed and placed in ice-cold carbogenated (95 % O_2_/5 % CO_2_) sucrose-modified artificial cerebrospinal fluid (in mM: 87 NaCl, 2.5 KCl, 25 NaHCO_3_, 1.25 NaH_2_PO_4_, 25 glucose, 75 sucrose, 7 MgCl_2_, 0.5 CaCl_2_). 400 μm thick thalamocortical (TC) slices were then cut on a Vibratome (VT1200s, Leica, Germany) and then stored submerged in sucrose-ACSF warmed to 34°C for 30 min and transferred to room temperature until needed.

### Whole-Cell Patch-Clamp Recordings

For electrophysiological recordings slices were transferred to a submerged recording chamber perfused with carbogenated normal ACSF (in mM: 125 NaCl, 2.5 KCl, 25 NaHCO_3_, 1.25 NaH_2_PO_4_, 25 glucose, 1 MgCl_2_, 2 CaCl_2_) maintained at near physiological temperatures (32 ± 1°C) with an inline heater (LinLab, Scientifica, UK) at a flow rate of 6-8 ml/min. Slices were visualized with IR-DIC illumination (BX-51, Olympus, Hamburg, Germany) initially with a 4x objective lens (N.A. 0.1) to position above a L4 barrel, and then with a 20x water-immersion objective (N.A. 1.0, Olympus). Whole-cell patch-clamp recordings were made with a Multiclamp 700B amplifier (Molecular Devices, USA). Recording pipettes were pulled from borosilicate glass capillaries (1.7 mm outer/1mm inner diameter, Harvard Apparatus, UK) on a horizontal electrode puller (P-97, Sutter Instruments, CA, USA), which when filled with intracellular solution gave a pipette resistance of 4-5 MΩ. Unless otherwise stated, all V-clamp recordings were performed at V_M_= -70 mV. All signals were filtered at 10 kHz using the built in 4-pole Bessel filter of the amplifier, digitized at 20 kHz on an analogue-digital interface (Digidata 1440, Axon Instruments, CA, USA), and acquired with pClamp software (pClamp 10, Axon Instruments, CA, USA). Data was analysed offline using the open source Stimfit software package (Guzman, Schlögl, and Schmidt-Hieber 2014) (http://www.stimfit.org). Cells were rejected if the I_hold_ was >150pA in voltage-clamp, membrane potential more depolarised than -50 mV in current-clamp, series resistance >30 MΩ, or the series resistance changed by more than 20% over the course of the recording.

### Sequential dendritic spine 2-photon glutamate uncaging

Slices were transferred to the recording chamber, which was perfused with normal ACSF, containing 50 µM picrotoxin (PTX) and 300 nM tetrodotoxin (TTX). For voltage clamp recordings of dendritic spine uncaging neurons were filled with an internal solution containing (in mM): 140 Cs-gluconate, 3 CsCl, 0.5 EGTA, 10 HEPES, 2 Mg-ATP, 2 Na_2_-ATP, 0.3 Na_2_-GTP, 1 phosphocreatine, 5 QX-314 chloride, 0.1% biotinoylated-lysine (Biocytin, Invitrogen, UK), and 0.1 AlexaFluor 488 or 594 (Invitrogen, UK), corrected to pH 7.4 with CsOH, Osm = 295 – 305 mOsm. Whole-cell patch clamp was then achieved and cells allowed to dye fill for 10 minutes prior to imaging. During this period, we collected 5 minutes of spontaneous recording, to analyse mEPSCs from recorded neurons at -70 mV voltage clamp. For all imaging and uncaging experiments we used a galvanometric scanning 2-photon microscope (Femto2D-Galvo, Femtonics, Budapest, Hungary) fitted with a femtosecond aligned, tuneable wavelength Ti:Sapphire laser (Chameleon, Coherent, CA, USA), controlled by a Pockel cell (Conoptics, CT, USA). Following dye filling, a short, low zoom z-stack was collected (2 µm steps, 2-3 pixel averaging, 512 x 512 pixels) over the whole dendritic extent of the cell at low laser power (<5 mW) with a high numerical aperture 20x lens (N.A. 1.0, Olympus, Japan). Then a short section of spiny dendrite, 50-100 µm from the cell somata and running parallel to the slice surface was selected and imaged at higher zoom. Between 7-10 spines were then selected based on being morphologically distinct from neighbouring spines, ordered distal to proximal to soma, and then 300 µM Rubi-Glutamate (Rubi-Glu; Ascent Scientific, Bristol, UK) was applied to the bath, and recirculated (total volume: 12.5 ml; flow rate: 6-8 mls/minute). Following wash-in of Rubi-Glu (<2 minutes), short duration, high power laser pulses (1 ms, λ780 nm, 80-100 mW, 0.2 µm diameter) local photolysis was performed ∼1 µm adjacent to individual spines. In a subset of recordings from WT mice, we confirmed spatial, quantal release, and pharmacological properties of Rubi-Glu uncaging under our recording conditions (Figure S1). Individual spines were sequentially uncaged at 2 second intervals followed by a 40 second pause; therefore each spine receiving Rubi-Glu photolysis every 60 seconds. All spines underwent photolysis at least 3 times and the average uncaging-EPSC (uEPSC) at -70 mV measured. In a subset of experiments we confirmed that these uEPSCs were mediated by direct activation of AMPARs by subsequent application of 10 µM CNQX to the perfusing ACSF (Figure S1D). Following each 3 repetition cycle, the focal plane and dendritic health was checked with short scans, at low power (<5 mW) to prevent background photolysis. Following successful recording of AMPA uEPSCs, we increased the holding potential to +40 mV and recorded the outward mixed AMPA/NMDA currents. In a subset of experiments we confirmed the AMPAR and NMDAR dependence of these outward currents by bath applying 10 µM CNQX and then 50 µM D-AP5 (Figure S1E). AMPA uEPSCs were measured over the first 10 ms following the uncaging stimulus (0.5 ms peak average) at both -70 and +40 mV. NMDA currents were measured from 20-50 ms post-photolysis, which was confirmed to be following complete decay of the AMPA uEPSC at -70 mV. All sequential spine uncaging experiments were performed as quickly as possible following dye filling, to prevent phototoxic damage to the recorded neurons, and L4 SCs resealed with an outside-out patch. Cells were rejected if photolysis resulted in blebbing of dendrites or depolarisation of the membrane potential.

In a subset of experiments, we performed mEPSC analysis of L4 SCs independent of Rubi-Glu photolysis, under the same conditions as above (with no AlexaFluor dye), recording 5 minutes of mEPSCs at -70 mV voltage clamp. Cells were then depolarised to +40 mV voltage-clamp and mixed AMPA/NMDA mEPSCs recorded for 1 minute, after which 10 µM CNQX was applied to the bath. Following full wash in of CNQX (∼2-3 minutes) a further 5 minutes of pure NMDA mEPSCs were recorded. In all experiments 50 µM APV was then bath applied, to confirm that the mEPSCs recorded were NMDAR-mediated. All mEPSC data was analysed using a moving-template algorithm (Clements and Bekkers 1997), with templates made from the tri-exponential non-linear fit to optimal mEPSCs at each holding potential using the event-detection interface of Stimfit. For mEPSCs at -70 mV, the minimum time between EPSCs was set to 7.5 ms, and 25 ms for those at +40 mV. Detected events were analysed if they had an amplitude greater than 3x the SD of the 5 ms preceding baseline of the mEPSC.

Summation of thalamic inputs to L4 SCs was measured by electrical stimulation of the ventrobasal thalamus with a twisted bipolar Ni-Chrome wire. Synaptically coupled barrels were identified by placing a field electrode (a patch electrode filled with ACSF) in visually identified barrels and stimulating the thalamus. When a field response was observed, then a L4 SC was recorded in whole-cell patch clamp, as described above. Trains of 5 stimuli were then delivered at 5-10 Hz, with a stimulation intensity sufficient to produce an EPSC of large amplitude similar between genotypes (20 to 540 pA; WT: 181 ± 35 pA; *Fmr1*^-/y^: 159 ± 34pA; D.F. = 23, t=0.44, P=0.66, T-test). In current clamp the EPSP summation was assessed as the ability of the recorded cell to fire an AP in response to this stimulus. Data are show as the average *P*_*spike*_ from 10 trials.

### Near-simultaneous dendritic spine 2-photon glutamate uncaging

To determine the summation properties of dendrites in L4 SCs we performed near simultaneous photolysis of Rubi-Glutamate at multiple dendritic spines (Branco, Clark, and Häusser 2010, Branco and Häusser 2011). Using a current-clamp optimized internal solution containing (in mM: 142 K-gluconate, 4 KCl, 0.5 EGTA, 10 HEPES, 2 MgCl_2_, 2 Na_2_-ATP, 0.3 Na_2_-GTP, 1 phosphocreatine, 0.1% Biocytin, and 0.1 AlexaFluor 488 (Invitrogen, UK), corrected to pH 7.4 with KOH, Osm = 295 – 305 mOsm) we dye filled neurons as for sequential photolysis described above, in normal ACSF containing PTX and TTX, but not Rubi-Glu. Once dye filling was complete (<10 minutes) we imaged the L4 SC (as above) at low zoom, then identified a superficial spiny dendrite 50-100 µm from the soma. At this point we placed a wide puff pipette (borosilicate patch pipette with tip broken to ∼20 µm diameter) just above the surface of the slice, adjacent to the dendrite of interest. The puff pipette was filled with 10 mM Rubi-Glu in a HEPES buffered ACSF (in mM: 140 NaCl, 2.5 KCl, 10 HEPES, 1.25 NaH_2_PO_4_, 25 glucose, 1 MgCl_2_, 2.5 CaCl_2_; adjusted to pH 7.4 with HCl). At this point the dendrite was imaged at high magnification and 7-10 spines chosen and a very low pressure stimulus given to the puff-pipette (3-5 mBar), sufficient to cause dialysis of the Rubi-Glu, but not powerful enough to cause obvious movement of the tissue. The dialysis of Rubi-Glu was maintained throughout the remainder of the recording. The cell was then switched to current-clamp mode, membrane potential held at -60 mV with a bias current, and spines 1-7 sequentially uncaged (0.5 ms laser duration, 80 mW power) to give the individual spines uEPSP amplitude. Following 3 repetitions and correction of focus, a line scan was created, with 0.5 ms dwell time at each spine ROI in order from distal to proximal. Spines were then uncaged in a cumulative manner, with 1, 2, 3 … n spines uncaged near simultaneously. The total duration of uncaging was 5.5 ms for 10 spines and there was a 10 second delay between each run of photolysis, with the total protocol lasting minimally 4-5 minutes. At least 3 repetitions of this protocol were run and focus re-checked. In a subset of experiments the HCN inhibitor ZD-7,288 (20 µM) was applied to the perfusing ACSF and a further 3 repetitions collected. All uEPSP data was analysed as peak amplitude measured over the 20 ms directly following beginning of the photolysis stimuli. Data was either normalised to the first EPSP amplitude, or measured as the absolute simultaneous uEPSP, as plotted against the summed individual uEPSP amplitude for the same spines.

In a set of experiments (without PTX, TTX or AlexaFluor 488), intrinsic electrophysiological properties of L4 SCs were measured, also in current-clamp mode. From resting membrane potential a hyper- to depolarizing family of current injections (−125 to +125 pA, 500ms duration) were given to the recorded neuron. The input resistance, rheobase current, and action potential discharge frequency were all measured from triplicate repetitions. In a further subset of experiments, 3x series of voltage steps were given (in voltage-clamp) from -60 mV to -110 mV (10 mV steps, 500 ms duration) to estimate the amplitude of I_h_ in the recorded L4 SCs. ZD-7,288 was then applied to the bath and the same steps repeated. I_h_ was estimated as the amplitude of the current produced in response to hyperpolarizing voltage steps.

### Visualisation and STED microscopy of recorded neurons

Following completion of experiments and resealing of the neuron, slices were immediately immersion fixed in 4% paraformaldehyde (PFA) overnight at 4 °C. Slices were then transferred to phosphate buffered saline (PBS; 0.025 M phosphate buffer + 0.9% NaCl; pH: 7.4) and kept at 4 °C until processed (<3 weeks). Slices were then cryoprotected in a solution containing 30% sucrose in PBS overnight at 4 °C and then freeze-thaw permeablised on lN_2_, and returned to cryoprotectant solution for 1 - 2 hrs. The slices were then mounted, recording side up, on the stage of a freezing microtome; which had been prepared with a plateau of OCT medium and slices embedded within OCT prior to sectioning. The OCT block containing the recorded slice was trimmed to the slice surface and then 50 µm sections taken from the top 200 µm. The sections were rinsed 3 times in PBS and then incubated with streptavidin conjugated to AlexaFluor488 (1:500, Invitrogen, UK) at 4 °C for 3-5 days. The slices were then washed for 2 hours in repeated washes of PBS and then desalted with PB and mounted on glass slides with fluorescence protecting mounting medium (Vectorshield, Vector Labs, UK).

Sections were imaged on a gated-Stimulated emission depletion (STED) microscope (SP8 gSTED, Leica, Germany). Cells were found using epifluorescent illumination (488 nm excitation) under direct optics at low magnification (20x air immersion objective lens, N.A. 0.75) and then positioned under high magnification (100x oil-immersion objective lens, N.A. 1.4, Olympus, Japan) and then switched to gSTED imaging. Sections were illuminated with 488 nm light, produced by a continuous-wave laser, and short sections of non-uncaged dendrite used to optimize acquisition parameters, first under conventional confocal detection, then by gSTED imaging. The 488 nm illumination laser was set to 60-70% of maximum power, and the continuous wave STED laser (592 nm) set to 25% and gated according to the best STED-depletion achievable in the samples (1.5 – 8 ms gating). Once optimized, a region of interest (ROI) was selected over the uncaged dendrite, which at 1024×1024 pixel size, gave a pixel resolution of 20-30 nm. Short stacks were taken over dendritic sections containing uncaged and non-spines (0.5 µm steps) with STED images interleaved with confocal images for confirmation of STED effect. STED images were deconvolved (Huygen’s STED option, Scientific Volume Imaging, Netherlands) and uncaged spines identified by comparison to live 2-photon images (see Figure 2A). Measurements of head width and neck length were then made on the deconvolved images in FIJI (ImageJ)(Schindelin et al. 2012).

### Serial block face scanning-electron microscopy (SBF-SEM) of L4 SCs

For SBF-SEM, 10 P14 mice (3 WT / 7 *Fmr1*^-/y^) were perfusion fixed. Briefly, mice were sedated with isoflurane and terminally anaesthetized with I.P. sodium pentobarbital (50 mg/mouse). The chest was opened and 10 mls of PBS (pH 7.4, filtered) transcardially perfused (∼0.5 mls/second); once cleared the PBS was replaced with ice-cold fixative solution containing (3.5% PFA, 0.5% glutaraldehyde, and 15% saturated picric acid; pH 7.4), and 20 mls perfused. Brains were then removed and post-fixed overnight at 4 °C in the same fixative solution. 60 µm coronal sections were cut on a vibratome (Leica VT1000) and S1 identified based on visual identification. Sections were then heavy-metal substituted: first sections were rinsed in chilled PBS (5 x 3 mins) and then incubated with 3% potassium ferrocyanide and 2% w/v OsO_4_ in PBS for 1 hr at 4 °C. Sections were rinsed liberally in double distilled (dd) H_2_O and then incubated with 1% w/v thiocarbohydroxide for 20 minutes at room temperature. Sections were rinsed again in ddH_2_0, and then incubated with 2% w/v OsO_4_ for 30 minutes at room temperature, rinsed in ddH_2_0 and contrasted in 1% w/v uranyl acetate overnight at 4 °C. Sections were rinsed in ddH_2_O and then contrasted with 0.6% w/v lead aspartate for 30 mins at 60 °C. Sections were then rinsed in ddH_2_O, dehydrated in serial dilutions of ethanol for 30 minutes each at 4 °C, then finally dehydrated twice in 100% ethanol and then 100% acetone both at 4 °C for 30 minutes. Sections were then impregnated with serial dilutions (25%, 50%, 75%, diluted in acetone) of Durcupan ACM (Sigma Aldrich, UK) at room temperature for 2 hours per dilution, followed by 100% Durcopan ACM overnight in a dissector at room temperature. Sections were transferred to fresh Durcupan ACM for 1 hour at room temperature and then flat-embedded on glass slides, coated with mould-release agent, cover-slipped, and then cured for 12 hours at 60 °C.

For SFB-SEM imaging, small pieces of L4 of S1 were dissected from flat-embedded sections, with aid of a stereo microscope and glued with cyanoacrylate to stage mounting pins. The mounted tissue was then trimmed and gold-plated prior to insertion imaging. Initially, semi-thin sections trimmed from the surface of the block, and imaged under transmission electron microscopy at low power to confirm tissue ultrastructure and ROI selection for SBF-SEM. Next the tissue blocks were mounted in an SBF-SEM (3View, Gatan, CA, USA) and 3 x ∼10 µm^2^ ROIs chosen on the surface of the block, avoiding blood vessels or L4 SC somata, and imaged at 50 nm steps at 8000x magnification (1024×1024, 10 nm pixel size). Approximately 100 sections were collected from each block, giving a total depth of 5 µm. SBF-SEM images were analysed offline using the TrakEM module of FIJI (Cardona et al. 2012). Dendrites and spines were traced as surface profiles and then PSDs identified on dendritic spines as electron dense regions within 25 nm of the lipid bilayer. 6-11 dendrites were reconstructed from each mouse, which possessed a total of 38-49 spines (average= 4.4 spines/dendrite). The incidence of PSDs was calculated as an average within each mouse, and final averages produced as an animal average.

### Data analysis

All data is presented as the mean ± SEM. All datasets were performed preliminarily (n = 3-4/genotype) and power analysis performed to determine group size required. Data examining the effect of single spine properties between genotype was analysed as a general linear mixed-effects model (GLMM), in which both animal and cell were chosen as random effects, and genotype and spine maintained as fixed effects. When animal average data is shown, datasets were tested for normality (d’Agostino-Pearson test) and either Student’s t-test or Mann-Whitney non-parametric U-test was performed. Comparison of linear regression was performed with a Sum-of-Squares F-test. Statistically significant differences were assumed if P<0.05. Which statistical test employed is indicated throughout the text. Either GraphPad Prism or R was used for all statistical analyses.

## Acknowledgements

The authors wish to thank: Drs. Alison Dunn and Rory Duncan of the Edinburgh Super-Resolution Imaging Consortium (ESRIC) for expert advice and technical support; Kathryn Whyte and Tracey Davey of the EM Research Service, Newcastle University, for technical assistance with electron microscopy. Funders: Simons Foundation Autism Research Initiative (529085), The Patrick Wild Centre, Medical Research Council UK (MR/P006213/1), BBSRC (BB/N015878/1), The Shirley Foundation and the RS Macdonald Charitable Trust.

## Author Contributions

SAB – designed and performed experiments, analysed/interpreted data and wrote the manuscript; APFD - designed and interpreted, performed experiments, analysed data and wrote the manuscript, ORD – analysed/interpreted data and wrote the manuscript, JTRI - designed experiments and wrote the manuscript, GEH - obtained funding, analysed/interpreted data and wrote the manuscript, DJAW – designed experiments, analysed/interpreted data, obtained funding and wrote the manuscript; PCK – designed experiments, analysed/interpreted data, obtained funding and wrote the manuscript

## Materials and Correspondence

All correspondence should be addressed to Peter C Kind

## References

Agmon, A, and BW Connors. 1991. “Thalamocortical responses of mouse somatosensory (barrel) cortexin vitro.” Neuroscience 41 (2):365–379.

Ashby, Michael C, and John TR Isaac. 2011. “Maturation of a recurrent excitatory neocortical circuit by experience-dependent unsilencing of newly formed dendritic spines.” Neuron 70 (3):510–521.

Bear, Mark F, Kimberly M Huber, and Stephen T Warren. 2004. “The mGluR theory of fragile X mental retardation.” Trends in neurosciences 27 (7):370–377.

Booker, Sam A, Jie Song, and Imre Vida. 2014. “Whole-cell patch-clamp recordings from morphologically-and neurochemically-identified hippocampal interneurons.” Journal of visualized experiments: JoVE (91).

Brager, Darrin H, Arvin R Akhavan, and Daniel Johnston. 2012. “Impaired dendritic expression and plasticity of h-channels in the fmr1−/y mouse model of fragile X syndrome.” Cell reports 1 (3):225–233.

Branco, Tiago, Beverley A Clark, and Michael Häusser. 2010. “Dendritic discrimination of temporal input sequences in cortical neurons.” Science 329 (5999):1671–1675.

Branco, Tiago, and Michael Häusser. 2011. “Synaptic integration gradients in single cortical pyramidal cell dendrites.” Neuron 69 (5):885–892.

Bureau, Ingrid, Gordon M. G. Shepherd, and Karel Svoboda. 2008. “Circuit and Plasticity Defects in the Developing Somatosensory Cortex of *Fmr1* Knock-Out Mice.” The Journal of Neuroscience 28 (20):5178–5188. doi: 10.1523/jneurosci.1076-08.2008.

Cardona, Albert, Stephan Saalfeld, Johannes Schindelin, Ignacio Arganda-Carreras, Stephan Preibisch, Mark Longair, Pavel Tomancak, Volker Hartenstein, and Rodney J Douglas. 2012. “TrakEM2 software for neural circuit reconstruction.” PloS one 7 (6):e38011.

Clements, JD, and JM Bekkers. 1997. “Detection of spontaneous synaptic events with an optimally scaled template.” Biophysical journal 73 (1):220–229.

Comery, Thomas A, Jennifer B Harris, Patrick J Willems, Ben A Oostra, Scott A Irwin, Ivan Jeanne Weiler, and William T Greenough. 1997. “Abnormal dendritic spines in fragile X knockout mice: maturation and pruning deficits.” Proceedings of the National Academy of Sciences 94 (10):5401–5404.

Contractor, Anis, Vitaly A Klyachko, and Carlos Portera-Cailliau. 2015. “Altered neuronal and circuit excitability in fragile X syndrome.” Neuron 87 (4):699–715.

Crocker-Buque, Alex, Sarah M Brown, Peter C Kind, John TR Isaac, and Michael I Daw. 2014. “Experience-dependent, layer-specific development of divergent thalamocortical connectivity.” Cerebral cortex 25 (8):2255–2266.

Darnell, Jennifer C, Sarah J Van Driesche, Chaolin Zhang, Ka Ying Sharon Hung, Aldo Mele, Claire E Fraser, Elizabeth F Stone, Cynthia Chen, John J Fak, and Sung Wook Chi. 2011. “FMRP stalls ribosomal translocation on mRNAs linked to synaptic function and autism.” Cell 146 (2):247–261.

Deng, Pan-Yue, David Sojka, and Vitaly A Klyachko. 2011. “Abnormal presynaptic short-term plasticity and information processing in a mouse model of fragile X syndrome.” Journal of Neuroscience 31 (30):10971–10982.

Fox, Kevin. 1992. “A critical period for experience-dependent synaptic plasticity in rat barrel cortex.” Journal of Neuroscience 12 (5):1826–1838.

Gibson, Jay R., Aundrea F. Bartley, Seth A. Hays, and Kimberly M. Huber. 2008. “Imbalance of Neocortical Excitation and Inhibition and Altered UP States Reflect Network Hyperexcitability in the Mouse Model of Fragile X Syndrome.” Journal of Neurophysiology 100 (5):2615–2626. doi: 10.1152/jn.90752.2008.

Gonçalves, J Tiago, James E Anstey, Peyman Golshani, and Carlos Portera-Cailliau. 2013. “Circuit level defects in the developing neocortex of Fragile X mice.” Nature neuroscience 16 (7):903.

Guzman, Segundo J, Alois Schlögl, and Christoph Schmidt-Hieber. 2014. “Stimfit: quantifying electrophysiological data with Python.” Frontiers in neuroinformatics 8.

Harlow, Emily G, Sally M Till, Theron A Russell, Lasani S Wijetunge, Peter Kind, and Anis Contractor. 2010. “Critical period plasticity is disrupted in the barrel cortex of FMR1 knockout mice.” Neuron 65 (3):385–398.

Harris, Kristen M, Frances E Jensen, and Beatrice Tsao. 1992. “Three-dimensional structure of dendritic spines and synapses in rat hippocampus (CA1) at postnatal day 15 and adult ages: implications for the maturation of synaptic physiology and long-term potentiation [published erratum appears in J Neurosci 1992 Aug; 12 (8): following table of contents].” Journal of Neuroscience 12 (7):2685–2705.

Irwin, Scott A., Roberto Galvez, and William T. Greenough. 2000. “Dendritic Spine Structural Anomalies in Fragile-X Mental Retardation Syndrome.” Cerebral Cortex 10 (10):1038–1044. doi: 10.1093/cercor/10.10.1038.

Jia, Hongbo, Zsuzsanna Varga, Bert Sakmann, and Arthur Konnerth. 2014. “Linear integration of spine Ca2+ signals in layer 4 cortical neurons in vivo.” Proceedings of the National Academy of Sciences 111 (25):9277–9282.

Lachiewicz, Ave M, Gail A Spiridigliozzi, Christina M Gullion, Sally N Ransford, and K Rao. 1994. “Aberrant behaviors of young boys with fragile X syndrome.” American Journal on Mental Retardation.

Lavzin, Maria, Sophia Rapoport, Alon Polsky, Liora Garion, and Jackie Schiller. 2012. “Nonlinear dendritic processing determines angular tuning of barrel cortex neurons in vivo.” Nature 490 (7420):397.

Losonczy, Attila, and Jeffrey C. Magee. 2006. “Integrative Properties of Radial Oblique Dendrites in Hippocampal CA1 Pyramidal Neurons.” Neuron 50 (2):291–307. doi: 10.1016/neuron.2006.03.016.

Magee, Jeffrey C. 1999. “Dendritic Ih normalizes temporal summation in hippocampal CA1 neurons.” Nature neuroscience 2 (6):508–514.

Magee, Jeffrey C, and Daniel Johnston. 1995. “Synaptic activation of voltage-gated channels in the dendrites of hippocampal pyramidal neurons.” Science 268 (5208):301.

Miller, LJ, DN McIntosh, J McGrath, V Shyu, M Lampe, AK Taylor, F Tassone, K Neitzel, T Stackhouse, and RJ Hagerman. 1999. “Electrodermal responses to sensory stimuli in individuals with fragile X syndrome.” Am J Med Genet 83:268–79.

Nikonenko, I., P. Jourdain, and D. Muller. 2003. “Presynaptic remodeling contributes to activity-dependent synaptogenesis.” Journal of Neuroscience 23 (24):8498–8505.

Petersen, Carl CH. 2007. “The functional organization of the barrel cortex.” Neuron 56 (2):339– 355.

Pfeiffer, Brad E, and Kimberly M Huber. 2007. “Fragile X mental retardation protein induces synapse loss through acute postsynaptic translational regulation.” The Journal of neuroscience 27 (12):3120–3130.

Pfeiffer, Brad E, and Kimberly M Huber. 2009. “The state of synapses in fragile X syndrome.” The Neuroscientist 15 (5):549–567.

Rall, Wilfrid. 1959. “Branching dendritic trees and motoneuron membrane resistivity.” Experimental Neurology 1 (5):491–527.

Schindelin, Johannes, Ignacio Arganda-Carreras, Erwin Frise, Verena Kaynig, Mark Longair, Tobias Pietzsch, Stephan Preibisch, Curtis Rueden, Stephan Saalfeld, and Benjamin Schmid. 2012. “Fiji: an open-source platform for biological-image analysis.” Nature methods 9 (7):676– 682.

Schubert, Dirk, Rolf Kötter, and Jochen F Staiger. 2007. “Mapping functional connectivity in barrel-related columns reveals layer-and cell type-specific microcircuits.” Brain Structure and Function 212 (2):107–119.

Till, Sally M, Lasani S Wijetunge, Viktoria G Seidel, Emily Harlow, Ann K Wright, Claudia Bagni, Anis Contractor, Thomas H Gillingwater, and Peter C Kind. 2012. “Altered maturation of the primary somatosensory cortex in a mouse model of fragile X syndrome.” Human molecular genetics:dds030.

Wang, Gordon X, Stephen J Smith, and Philippe Mourrain. 2014. “Fmr1 KO and Fenobam Treatment Differentially Impact Distinct Synapse Populations of Mouse Neocortex.” Neuron 84 (6):1273–1286.

White, Edward L, and Michael P Rock. 1980. “Three-dimensional aspects and synaptic relationships of a Golgi-impregnated spiny stellate cell reconstructed from serial thin sections.” Journal of neurocytology 9 (5):615–636.

Wijetunge, Lasani S., Julie Angibaud, Andreas Frick, Peter C. Kind, and U. Valentin Nägerl. 2014. “Stimulated Emission Depletion (STED) Microscopy Reveals Nanoscale Defects in the Developmental Trajectory of Dendritic Spine Morphogenesis in a Mouse Model of Fragile X Syndrome.” The Journal of Neuroscience 34 (18):6405–6412. doi: 10.1523/jneurosci.5302-13.2014.

Zhang, Yu, Audrey Bonnan, Guillaume Bony, Isabelle Ferezou, Susanna Pietropaolo, Melanie Ginger, Nathalie Sans, Jean Rossier, Ben Oostra, Gwen LeMasson, and Andreas Frick. 2014. “Dendritic channelopathies contribute to neocortical and sensory hyperexcitability in Fmr1-/y mice.” Nat Neurosci 17 (12):1701–1709. doi: 10.1038/nn.3864.

Zoghbi, Huda Y, and Mark F Bear. 2012. “Synaptic dysfunction in neurodevelopmental disorders associated with autism and intellectual disabilities.” Cold Spring Harbor perspectives in biology:a009886.

